# Generality and opponency of rostromedial tegmental (RMTg) roles in valence processing

**DOI:** 10.1101/410308

**Authors:** Hao Li, Dominika Pullmann, Jennifer Y. Cho, Maya Eid, Thomas C. Jhou

## Abstract

The rostromedial tegmental nucleus (RMTg), a GABAergic afferent to midbrain dopamine (DA) neurons, has been hypothesized to encode aversive stimuli. However, this encoding pattern has only been demonstrated for a limited number of stimuli, and its influence on the ventral tegmental (VTA) responses to aversive stimuli is untested. Here, we found that RMTg neurons show average inhibitions to rewarding stimuli and excitations to aversive stimuli of greatly varying sensory modalities and timescales. Notably, negative valence-encoding neurons are particularly enriched in subpopulations projecting to the VTA versus other targets. Additionally, RMTg neurons also dynamically encode “opponent” changes in motivational states induced by removal of sustained stimuli. Finally, excitotoxic RMTg lesions impair conditioned place aversion to multiple aversive stimuli, and greatly reduce aversive stimulus-induced inhibitions in VTA neurons, particularly in putative DA-like neurons. Together, our findings indicate a broad RMTg role in encoding aversion and potentially driving DA responses and behavior.

## Introduction

The rostromedial tegmental nucleus (RMTg), or tail of the ventral tegmental area, is a GABAergic nucleus first described in 2009 as one of the major inhibitory inputs to neurons in the ventral tegmental area (VTA) (Jhou et al., 2009; Kaufling et al., 2009). Previous studies have shown that RMTg axons overwhelmingly synapse on tyrosine hydroxylase (TH) positive neurons in the VTA and substantia nigra, and electrical stimulation of the RMTg dramatically suppresses DA neuron firing (Balcita-Pedicino et al., 2011; Bourdy et al., 2014). In the past decade, increasing numbers of studies have suggested an important RMTg role in conveying information about aversive stimuli onto VTA DA neurons and mediating behavioral responses to these aversive stimuli (Brown et al., 2017; Hong et al., 2011; Jhou et al., 2009; Jhou et al., 2013; Vento et al., 2017). However, some fundamental questions about the proposed RMTg role in aversive encoding have not been addressed.

Most prior studies had only tested RMTg responses to a limited range of aversive stimuli, e.g. footshocks and airpuffs (Hong et al., 2011; Jhou et al., 2009). One notable exception is a recent study that examined 11 distinct aversive stimuli, but found that only 3 of them increased RMTg expression of the immediate-early gene c-Fos (Sanchez-Catalan et al., 2017). Because c-Fos is often used as a proxy for neuronal firing, this result raises questions about the generalizability of RMTg responses to other aversive stimuli, albeit with the caveat that c-Fos likely reflects intracellular signaling events rather than firing per se (Kovacs, 2008).

In addition to the uncertainty in generalizability to aversive stimuli, earlier studies had also failed to characterize the influence of RMTg neurons on DA responses to aversive stimuli, raising further questions about the proposed RMTg role. Increasingly many studies have noted that DA responses to aversive stimuli, although somewhat complex, often exhibit a prominent inhibitory component (Fiorillo et al., 2013; Matsumoto et al., 2016; Tian and Uchida, 2015), and that the aversive stimulus-induced inhibition is associated with behavioral avoidance and diminished learning (Chang et al., 2016; Lammel et al., 2012). Previous study has suggested that these inhibitory responses are primarily driven by GABAergic transmissions onto DA neurons (Henny et al., 2012). However, the sources of the GABAergic transmissions are still unknown, and have been proposed to arise from numerous possible sources including not only the RMTg, but also the lateral hypothalamus, ventral pallidum, and extended amygdala (Jennings et al., 2013; Tian et al., 2016).

To address these questions about generalizability of RMTg responses to aversive stimuli and its influence on DA neurons, we recorded VTA and RMTg neuron responses to a wide range of aversive stimuli in freely-moving rats, while using excitotoxic lesions to investigate the RMTg influence on VTA neuron firings and on conditioned place aversion to these stimuli.

## Results

### RMTg neurons are activated by diverse phasic aversive stimuli

To examine the generalizability of RMTg responses to aversive stimuli, we recorded RMTg neuron responses to six distinct aversive stimuli covering a broad range of stimulus modalities. Three of these stimuli were phasic: footshock (0.7 mA), loud siren (115dB), and bright light (1600 lumens), which lasted for 10ms, 1 second, and 2 seconds respectively. Three additional stimuli were sustained: lithium chloride (LiCl), restraint stress, and aversive effect of cocaine, lasting for several minutes in duration (**Fig. 1A**). Stimuli were chosen to represent distinct sensory modalities, whose aversive qualities had been noted previously (Ettenberg et al., 1999; Tzschentke, 2007; Winston et al., 2001). All recorded neurons were also tested for their response to reward cues and neutral tones (75dB) that were followed by a sucrose pellet delivery and no consequence, respectively.

**Figure 1.**
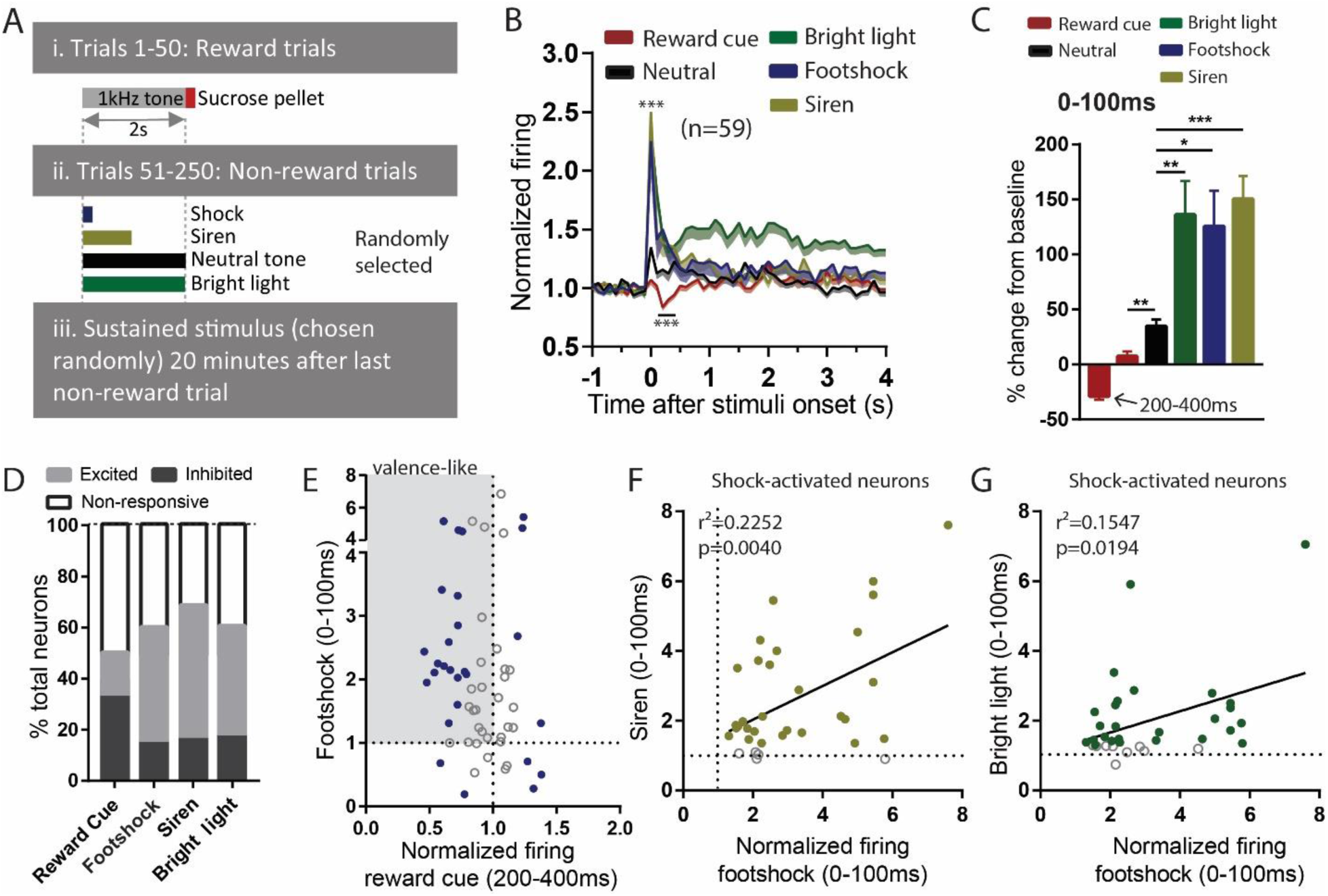
RMTg neurons are activated by diverse phasic aversive stimuli. (**A**) Schematic of recording paradigm. (**B, C**) RMTg neurons on average showed inhibition to reward-predictive cues during 200-400ms post-stimulus window, and small excitations to neutral tones and large excitations to footshock, siren, and bright light during 0-100ms post-stimulus windows. (**D**) Percentage of RMTg neurons that showed inhibition, excitation or no responses to stimuli (200-400ms window for reward cues, and 0-100ms window for rest stimuli). (**E**) Scatterplot of individual neurons’ responses to reward cues and footshocks. Many reward-cue inhibited neurons were also excited by footshocks, consistent with valence-encoding pattern. Blue solid dots: neurons significantly responded to both reward cues and footshocks. Gray shaded box: valence-consistent quadrant (neurons were inhibited by reward cues and excited by footshocks.) (**F, G**) RMTg neurons activated by footshocks tended to also be activated by siren and bright light, in proportion to the magnitude of response to footshock. Solid dots: neurons significantly responded to siren and bright light.

Out of 151 recorded neurons, 59 were located in the RMTg, as defined by immunostaining for FOXP1 (Lahti et al., 2016) (**Supplementary Fig. 1A, B**). On average, RMTg neurons showed significant inhibition to reward cues during 200-400ms post-stimulus (z=-5.03), and rapid excitations to all other phasic stimuli during 0-100ms post-stimulus (z=5.76, z=3.87, z=7.18, and z=4.44 for neutral tone, footshock, siren, and bright light, respectively) (**Fig. 1B**). Moreover, responses to the three phasic aversive stimuli (footshock, siren, and bright light) were significantly greater than responses to the neutral tone, suggesting an aversion-related augmentation in RMTg responses to these stimuli (F=1.582, p=0.027, p=0.0001, and p=0.0048 for neutral tone compared with footshock, siren, and bright light, repeated measures one-way ANOVA) (**Fig. 1C**).

When individual neurons (instead of population averages) were analyzed, we observed considerable variation among individual neuron responses to these stimuli. Specifically, we found 51% (30/59) of RMTg neurons showing significant changes relative to baseline firing (in either direction) to reward cue, with 2 neurons showing ramping activities, and almost three-fourths of these responsive neurons (22/30=73%) being inhibited largely during a window 200-400ms post-stimulus. Furthermore, 59% (35/59), 70% (41/59), and 61% (36/59) of RMTg neurons showed significant responses (in either direction) to footshock, siren, and bright light, again with roughly three-fourths showed excitations (27/35 = 77%, 31/41 = 75%, and 26/36 = 72% for footshock, siren, and light, respectively, 0-100ms post-stimulus) (**Fig. 1D**). Interestingly, we noticed that almost all (18/22) reward-cue inhibited neurons showed significant excitations to footshock, even though shock-excited neurons are only 31% of all RMTg neurons, suggesting a negative valence-encoding pattern specifically in the reward-cue inhibited population (**Fig. 1E**). Moreover, among RMTg neurons significantly excited by footshock, these responses were positively correlated with responses of each individual neuron to siren and bright light, i.e. neurons more strongly activated by the footshock were also more strongly activated by siren and bright light (r^2^=0.2252, p=0.004 and r^2^=0.1547, p=0.0194 for siren and light respectively, 0-100ms post-stimulus) (**Fig. 1F, G**).

### RMTg neurons exhibit biphasic responses to sustained aversive stimuli consistent with opponent process theory

In addition to testing RMTg responses to multiple phasic aversive stimuli, we also examined these same neurons’ responses to one of several sustained aversive stimuli lasting several minutes each. These were: a low dose of LiCl (10 mg/kg i.p.), restraint stress (6 minutes), or cocaine (0.75 mg/kg i.v.), which produces an aversive “crash” several minutes after an initial rewarding phase. All stimuli were administered roughly 20 minutes after rats had been tested with phasic stimuli (**Fig. 1A**), allowing subsequent comparison of neural responses across multiple stimuli. Notably, the dose of LiCl that we used is relatively low compared to the much higher doses commonly used to induce conditioned taste aversions, and is thought to produce modest aversive effects lasting only approximately 15 minutes (Tomasiewicz et al., 2006).

All of these sustained stimuli produced two distinct phases of response in RMTg firing. We found that both LiCl (n=22) and restraint stress (n=12) increased RMTg firing for several minutes after the onset of each stimulus (first 0-10 minutes and 0-3 minutes, respectively) (p=0.01 and p=0.02, one-way ANOVA). These periods of activation are somewhat shorter than the expected duration of the aversion for each stimulus (15 and 6 minutes, respectively), suggesting some habituation of the RMTg excitation even while the stimulus is still ongoinog. Interestingly, RMTg firing was reduced 20 minutes after LiCl injection or shortly after cessation of restraint stress, exhibiting a “rebound” inhibition to below baseline levels (20-30 minutes post-stimulus for LiCl and 9-12 minutes for restraint stress, p=0.047, and p=0.036, respectively, one-way ANOVA) (**Fig. 2A, E**). In contrast, saline injections of equal volume as the LiCl injections had no effect on RMTg firing during either of the two time windows where LiCl had produced responses (n=12, p=0.23 and p=0.84, respectively, one-way ANOVA) (**Fig. 2A**). As with phasic stimuli, we saw heterogeneous responses to LiCl and restraint stress. Roughly 36-42% RMTg neurons showed both an initial excitation and a rebound inhibition, again suggesting a negative valence-encoding pattern (8/22 and 5/12 neurons for LiCl and restraint stress, respectively) (**Fig. 2B, F**). The remaining neurons typically showed responses during only one of the two phases. We conducted further analyses to examine whether responses during initial and rebound phases were consistent with encoding of motivational states. We found that RMTg responses during the initial phase of LiCl and restraint stress were positively correlated with responses of these same neurons to footshock (r^2^=0.1469, p=0.0033 and r^2^=0.3783, p=0.033, respectively) (**Fig. 2C, G**), while their rebound phase responses correlated with responses to reward cues (r^2^=0.2165, p<0.0001 and r^2^=0.3954, p=0.038, respectively) (**Fig. 2D, H**).

**Figure 2.**
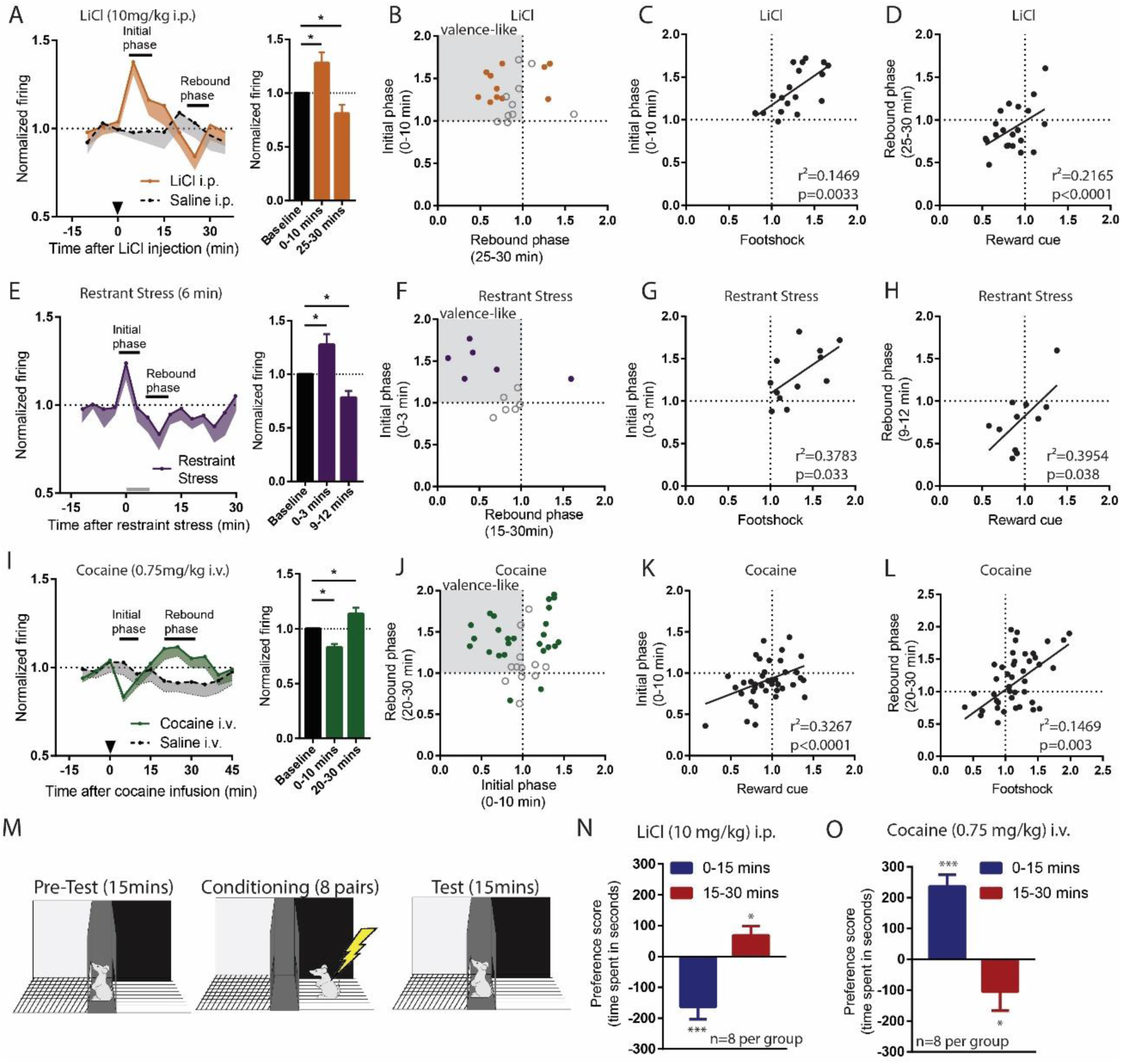
RMTg neurons exhibit biphasic responses to sustained aversive stimuli consistent with opponent process theory. (**A, B**) RMTg neurons showed activation to a low dose of LiCl for 10 minutes post injection, and (**E, F**) to restraint stress during the first 3 minutes of the 6-min restraint. In both cases, aversive stimulus also resulted in a rebound inhibition of firing below baseline during a later time window (20-30 minute window for LiCl, and 9-12 minute window for restraint stress). (**C, D**) Individual responses during initial phases of LiCl and restraint stress were correlated with their responses to footshock, while responses during rebound phases were correlated with their responses to reward cue (**G, H**). (**I, J**) Cocaine infusion inhibited RMTg neurons during the first 10 minutes and excited RMTg neurons during 15-25 minutes post infusion. (**K, L**) Individual responses during rebound phase of cocaine were correlated with their responses to footshock, while responses during initial phase were correlated with their responses to reward cue. (**M**) Schematic of conditioned place preference regimen. (**N**) Low dose of LiCl (10mg/kg) i.p. injection induced place aversion during 0-15 minutes and place preference during 15-30 minutes post-stimulus. (**O**) Cocaine (0.75 mg/kg) i.v. infusion induced place preference during 0-15 minutes and place aversion 15-30 minutes post-stimulus. Solid dots in **B**, **F**, and **J**: neurons significantly responded during both initial and rebound phases.

As LiCl and restraint stress are both aversive, we hypothesized that the rebound inhibitions of RMTg neurons may be correlated with the rewarding “relief” upon the removal of these stimuli. Conversely, we hypothesized that the converse might also be true, that RMTg neurons might show a rebound *excitation* after removal of a sustained rewarding stimulus, as we had observed earlier in the LHb (Jhou et al., 2013). Thus, we further examined RMTg responses to a 0.75 mg/kg i.v. cocaine infusion, which previous studies had shown produce rewarding effects for about 10 minutes followed by an aversive “crash” beginning around 15 minutes post-injection (Ettenberg et al., 1999; Jhou et al., 2013). We found that RMTg neurons (n=38) showed the similar bi-phasic responses to cocaine as to LiCl and restraint stress, but in the opposite direction. Specifically, they were initially inhibited by cocaine 0-10 minutes post-infusion (initial phase), and subsequently activated 20-30 minutes post-infusion (rebound phase), when cocaine was aversive (p=0.0001 and p=0.038, one-way ANOVA). Saline infusions (n=12) had no effect on RMTg firing during either phase (p=0.62 and p=0.25, one-way ANOVA) (**Fig. 2I**). Furthermore, when individual neuron responses were analyzed, we found that 32% of RMTg neurons (12/38) exhibited both inhibition during the initial phase and excitation during the rebound phase, again consistent with the negative valence-encoding pattern seen previously (**Fig. 2J**). Furthermore, responses during the initial phase were positively correlated with responses of these same neurons to food-predictive cues (r^2^=0.3267, p<0.0001), while responses during the rebound phase were positively correlated with responses to footshocks (r^2^=0.1469, p=0.003).

Although these results imply that RMTg neurons bi-directionally encode aversive and rewarding properties of sustained stimuli, we did not directly test whether the early and rebound phases of LiCl and cocaine are indeed aversive and rewarding. Thus, we tested four groups of animals (8 per group) for conditioned place preference or aversion to the same doses of LiCl and cocaine as during the recordings. We placed animals into conditioning chambers either 0-15 minutes or 15-30 minutes post-stimulus, to approximately match neural responses to initial and rebound phases seen in earlier recordings (**Fig. 2M**). In separate groups of LiCl-treated groups, we found that animals developed place aversion when placed into conditioning chambers immediately after the injection (z=-3.98), but showed place preference when placed into chambers 15 minutes after the injection (z=2.29) (**Fig. 2N**). In contrast, separate groups of cocaine-treated animals developed place preference if conditioned immediately after infusion (z=-1.78), and place aversion if conditioned 15 minutes later (z=6.31) (**Fig. 2O**). Together, our results indicate that bidirectional behavioral effects of sustained rewarding or aversive stimuli correlate with bidirectional responses of RMTg neurons to these stimuli, suggesting a possible substrate for an “opponent” responses to stimuli described decades ago by opponent process theory (Solomon and Corbit, 1973).

### VTA-projecting RMTg neurons preferentially show valence-encoding patterns

As previous results have indicated heterogeneities in RMTg responses to both phasic and sustained stimuli, with only 30%-40% of RMTg neurons encoding negative valence, we next examined whether the heterogeneity in response patterns might be related to heterogeneity in RMTg projection targets. Thus, we used endoscopic calcium imaging and selectively recorded RMTg neurons that project to either the VTA or the dorsal raphe nucleus (DRN), regions that are both related to encoding of motivational stimuli (Cohen et al., 2012; Li et al., 2016). Because of the need to keep GRIN lenses short, this experiment was performed in mice, rather than rats, but the anatomy of the RMTg is very similar between species (R.S. Smith and T.C. Jhou, unpublished data). We injected into wild type mice a retrogradely transported canine adenovirus expressing Cre recombinase (CAV2-Cre) into either the VTA or (in separate mice) the DRN, along with a second virus into the RMTg expressing a Cre-dependent fluorescent calcium indicator (gCaMP6f) (**Fig. 3A-C, Supplementary Fig. 2A, B**). Mice were tested in separate sessions in which they received either a footshock (0.3mA), or an auditory tone followed by food pellet delivery.

**Figure 3.**
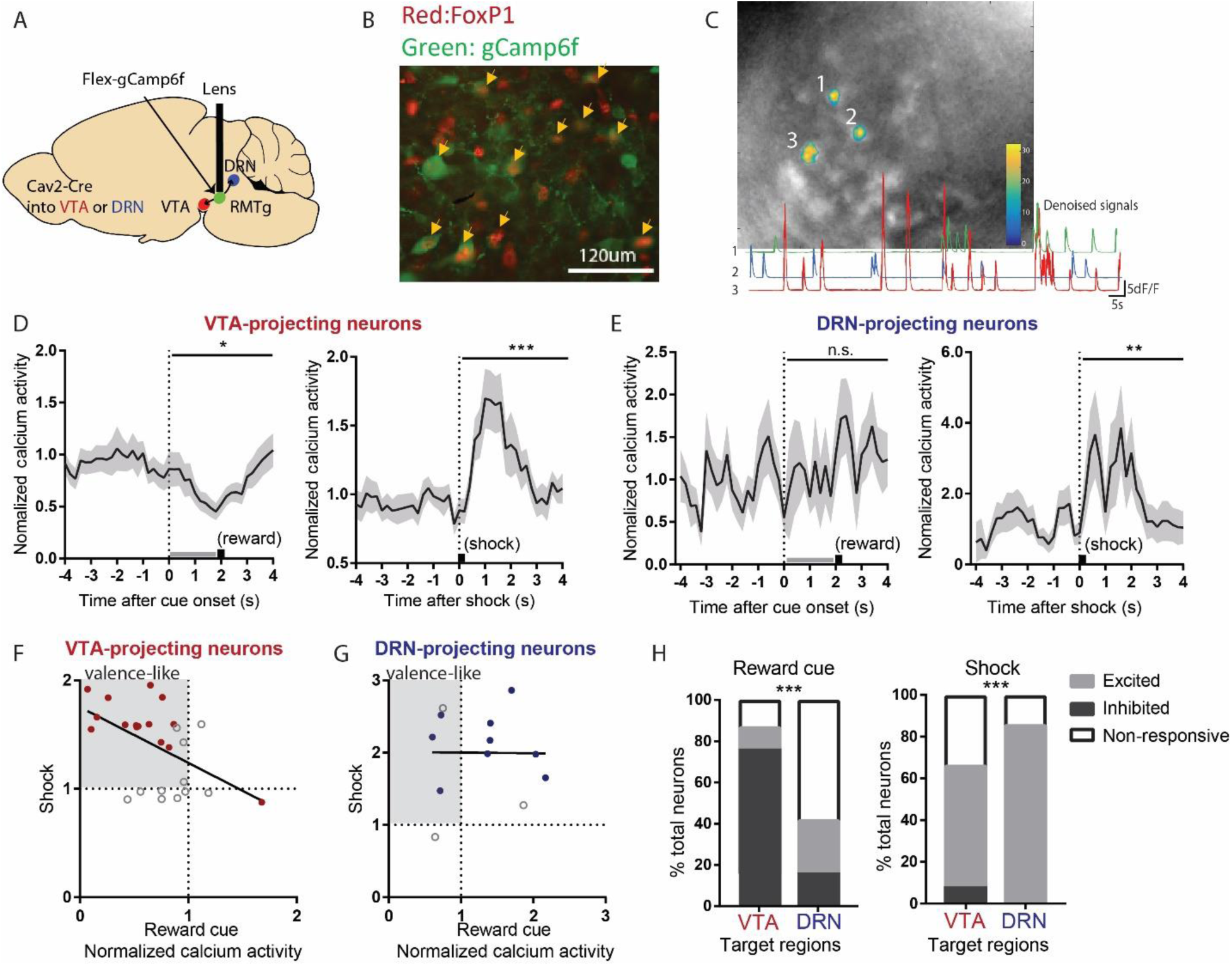
VTA-projecting RMTg neurons preferentially show valence-encoding patterns. (**A**) We injected a Cav2-cre into the VTA or DRN, which was retrogradely transported to the RMTg in mice, driving gCaMP6f expression in subsets of RMTg neurons projecting to the VTA or DRN. (**B**) Representative photograph of the RMTg region in which gCaMP6 in RMTg (green label) is co-expressed with FOXP1 (red), a transcription factor locally specific to RMTg neurons. (**C**) A representative photo of gCaMP6f positive neurons *in vivo* (upper panel) and denoised Ca^2+^ traces extracted from the marked neurons (lower panel). (**D**) VTA-projecting RMTg neurons showed an average inhibition by reward cues, and excitation by footshocks. (**E**) DRN-projecting neurons showed no average response to reward cues, but were excited by footshocks. (**F**) Among VTA-projecting RMTg neurons, neurons showing stronger excitations to shock tended to also show stronger inhibitions to the reward cue, while individual DRN neurons did not show this correlation (**G**). Colored solid dots: neurons significantly responded to both reward cues and footshocks. (**H**) VTA-projection neurons were much more likely to be inhibited by the reward cue than DRN-projecting neurons, while VTA- and DRN-projecting neurons were both predominantly activated by shock, but with a slightly higher probability for DRN-projecting neurons.

We found that VTA-projecting neurons were on average inhibited by reward cues and activated by footshocks, i.e. a negative valence-encoding pattern (**Fig. 3D, Supplementary Fig. 2C, D**). Analysis of individual neurons further confirmed this; among 25 VTA-projecting neurons, a majority (14/25 neurons) were inhibited by reward cues and activated by footshocks, while only 7/25 neurons responded to only one of the two stimuli, with the remaining 4 neurons showing no response to either stimulus. Furthermore, their responses to the reward cues correlated negatively with responses to footshocks (r^2^=0.2661, p=0.0070) (**Fig. 3F**).

In marked contrast to VTA-projecting neurons, DRN-projecting neurons on average showed no responses to reward cues, although there was an average activation by footshocks, similar to VTA-projecting neurons (**Fig. 3E**). Analysis of individual neurons further showed that responses to reward cues and footshocks did not correlate with each other (p=0.9775) (**Fig. 3G**). Overall, VTA- and DRN-projecting neurons exhibited markedly different proportions of neurons that were activated versus inhibited by reward cues or footshocks (p<0.001 for both footshock and reward cue, Chi-square) (**Fig. 3H**). Thus, our results indicate that VTA-projecting but not DRN-projecting RMTg neurons appear highly enriched in negative valence-encoding patterns.

### Excitotoxic RMTg lesions abolished VTA inhibitions by aversive stimuli

Our findings that RMTg neurons are activated by aversive stimuli and VTA-projecting neurons are preferentially negative valence-encoding suggest that the RMTg could drive VTA responses to aversive stimuli, a hypothesis we tested by recording VTA neuron responses to phasic aversive stimuli with or without RMTg lesions (**Fig. 4E, Supplementary Fig. 1C, D**). NeuN staining showed that lesions were mostly restricted in the RMTg area and did not extend to surrounding structures such as pedunculopontine nucleus (PPTg) and dorsal raphe nucleus (DRN) (**Fig. 4E**). We classified recorded VTA neurons as being putative dopamine neurons (pDA neurons) by their phasic activations to reward cues (Sham: n=21 pDA, n=45 non-pDA, Lesion: n=17 pDA, n=40 non-pDA), an activity signature repeatedly shown to correlate highly with optogenetically identified DA neurons in mice (Cohen et al., 2012; Eshel et al., 2015; Matsumoto et al., 2016) (**Fig. 4A, B**). We also observed about 1/3 of recorded VTA neurons in sham group exhibiting ramping activities in responses to reward cues (putative GABA neurons or pGABA), which are consistent with firing patterns found in genetically identified GABAergic neurons in the VTA (Cohen et al., 2012) (**Fig. 4B**). Additionally, 4 of all recorded VTA neurons in sham group showed negative valence encoding, predominantly seen in the RMTg.

**Figure 4.**
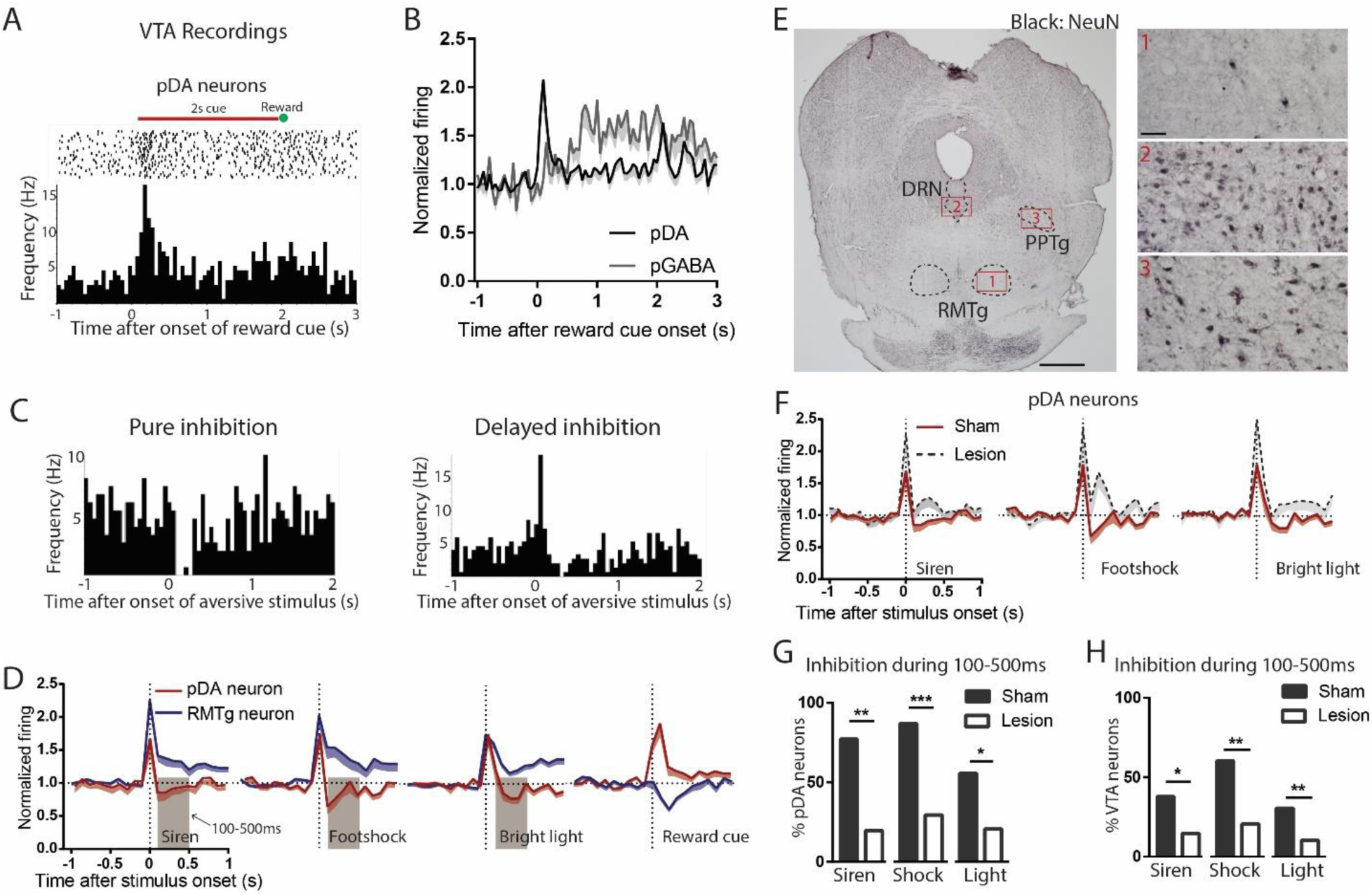
Excitotoxic RMTg lesions abolished VTA inhibitions by aversive stimuli. (**A, B**) pDA neurons in the VTA were classified by their phasic activation to reward cues. (**C**) Raster plots of pure inhibition and delayed inhibition response types observed in pDA neuron after aversive stimuli. (**D**) Comparisons of RMTg (blue trace) and pDA (red trace) neuron responses to affective stimuli. All three aversive stimuli elicited initial excitations in both RMTg and pDA neurons, after which RMTg neurons remained excited while pDA neurons showed inhibition during 100-500ms window post-stimulus (brown-shaded boxs). pDA neuron activation to reward cues was much faster than RMTg inhibition to the same cue, making it unlikely that responses to the reward cue would be driven by the RMTg. (**E**) NeuN staining showed that lesions were mostly restricted in the RMTg area and did not extend to surrounding structures such as pedunculopontine nucleus (PPTg) and dorsal raphe nucleus (DRN). Scalebars: 1mm and 100um for left and right panels. (**F**) RMTg lesion (dashed trace) eliminated aversion-induced inhibition in pDA neurons. (**G**) Bar graphs again showing loss of aversion-induced inhibition during 100-500ms in pDA neurons after RMTg lesions. (**H**) Loss of aversion-induced inhibition in all recorded VTA neurons.

We found that most pDA neurons in the sham group showed inhibitions to aversive stimuli, in some cases after a brief initial excitation (**Fig. 4C**). Specifically, 86% of stimulus-responsive pDA neurons were inhibited by footshock (54% showing only inhibition; 32% showing delayed inhibition which occurred after a brief initial excitation), 75% by siren (28% showing only inhibition; 47% showing delayed inhibition), and 56% by bright light (27% showing only inhibition; 29% showing delayed inhibition), while the remaining showed excitation only (**Supplementary Fig. 3A-C**). Notably, inhibition by aversive stimuli was most prominent during a 100-500ms window post-stimulus, while activation by reward cues were most prominent in a 0-200ms window post-stimulus, a pattern opposite to that of RMTg neurons in which responses to aversive stimuli tended to be faster (**Fig. 4D**).

Among recorded VTA neurons, we found that RMTg lesions did not affected the magnitude or the percentage of neurons responding to reward cues (0-200ms post-stimulus) (p>0.05, two-way ANOVA, p=0.879, Chi-square) (**Supplementary Fig. 3D, E**). Hence, we further analyzed reward-activated VTA neurons on the presumption that these were pDA in both lesioned and unlesioned animals. We found that, after RMTg lesions, pDA neurons no longer showed an average inhibition to any of the three phasic aversive stimuli (**Fig. 4F**), and the proportions of pDA neurons inhibited by aversive stimuli were dramatically reduced to 19%, 16%, and 18% for footshock, siren, and bright light, respectively (p<0.0001, p=0.008, and p=0.048, Chi-square, 100-500ms post-stimulus) (**Fig. 4G**). Due to the potential inaccuracy of our classification of pDA neurons, we then analyzed all recorded VTA neurons instead of pDA neurons alone. Consistent with pDA analysis, the proportions of inhibitory responses were also reduced by RMTg lesions, although to a less extreme degree than the pDA population. Specifically, in intact animals, 59%, 42%, and 36% of VTA neurons showed significant inhibitions to footshock, siren, and bright light, while in RMTg-lesioned rats, these proportions were reduced to 21%, 16%, and 9%, respectively (p=0.005, p=0.017, and p=0.004, for footshock, siren, and bright light, respectively, Chi-square) (**Fig. 4H**).

In addition to the reductions in inhibitory responses to aversive stimuli, we observed non-significant trends towards increases in proportions of pDA neurons excited by these stimuli (p=0.182, p=0.047, and p=0.766 for siren, footshock, and bright light, respectively, chi-square) (**Supplementary Fig. 3G**). As noted above, many pDA neurons exhibit initial excitations to aversive stimuli, often preceding subsequent inhibitions. RMTg lesions also increased the magnitudes of these initial excitations relative to shams (p<0.05, unpaired t-test), and caused non-significant trends towards increases in proportions of pDA neurons exhibiting these initial excitations (p>0.05, chi-square) (**Supplementary Fig. 3H**). Notably, RMTg lesions did not alter basal firing rates of pDA neurons (5.4 Hz vs 5.2 Hz, n=23, p=0.507, unpaired t-test). However, we did find that RMTg lesions increased the percentage of spikes found in bursts of pDA neurons (p = 0.014, unpaired t-test) (**Supplementary Fig. 3F**). Together, RMTg lesions influence both inhibitions and excitations of pDA neurons to aversive stimuli, greatly reducing inhibitions, and modestly increasing excitations to these stimuli.

### RMTg lesions disrupt conditioned place aversion to a wide range of stimuli

Finally, we examined the effect of RMTg lesions on conditioned place test for three of the aversive stimuli that we tested earlier: siren, bright light, and LiCl (**Fig. 5A, Supplementary Fig. 1C**). Using a three-chambered apparatus, we were able to measure the effect of conditioning on both the time spent in each chamber (stimulus-paired, unpaired, and neutral) as well as the relative number of entries into the paired and unpaired chambers (**Fig. 2M**). We found that on average, sham rats expressed a significant aversion to the light and LiCl, as measured by both time spent in, and entries into, the stimulus-paired versus unpaired chambers (p= 0.012 and p<0.0001, respectively for time, and p=0.033 and p<0.0001 for entries, one-tailed t-test). Shams also expressed a significant aversion to the siren as measured by relative number of entries into the paired versus unpaired chamber. However, this aversion did not reach significance as measured by total time spent (p=0.003 and p=0.067, respectively for entries and time, one-tailed t-test). A two-way ANOVA comparing lesioned versus sham groups then showed a significant main effect of lesion for both time and entries (p=0.003 each), indicating that lesioned rats showed smaller (or in some cases reversed) biases away from the stimulus-paired chambers. When examining specific stimuli, the lesion effect only reached significance for the light, but not LiCl and siren (p=0.728, p=0.036, and p=0.209 for siren, bright light and LiCl, respectively, corrected for multiple comparisons) (**Fig. 5B**). RMTg lesions produced a somewhat stronger effect on the preference score as measured by entries, with the two-way ANOVA again showing a main effect of lesion (p<0.0001), with this effect also being significant for each stimulus (p=0.017, p=0.003, and p=0.017 for siren, LiCl, and bright light, respectively) (**Fig. 5C**). Although preference scores could be confounded if RMTg lesions produce extreme motoric disinhibition, we found this was not the case; in particular, lesion group showed slightly greater overall locomotor activities measured by photobeam counts (p=0.02, unpaired t-test) (**Fig. 5D**), but lesion and sham groups did not show differences in total chamber entries prior to conditioning, suggesting that effects of RMTg lesions on locomotion were minor (p=0.106, unpaired t-test) (**Fig. 5E**). Interestingly, aversive conditioning reduced total chamber entries for all three stimuli in the sham group (p=0.04, p=0.0004, and p=0.001 for siren, bright light and LiCl, respectively, paired t-test) (**Fig. 5F**), but not the RMTg lesioned group (p=0.34, p=0.22, and p=0.01 for siren, bright light and LiCl, respectively, paired t-test) (**Fig. 5G**), suggesting an overall inhibitory effect of aversive conditioning in normal rats that was not present in the lesioned group.

**Figure 5.**
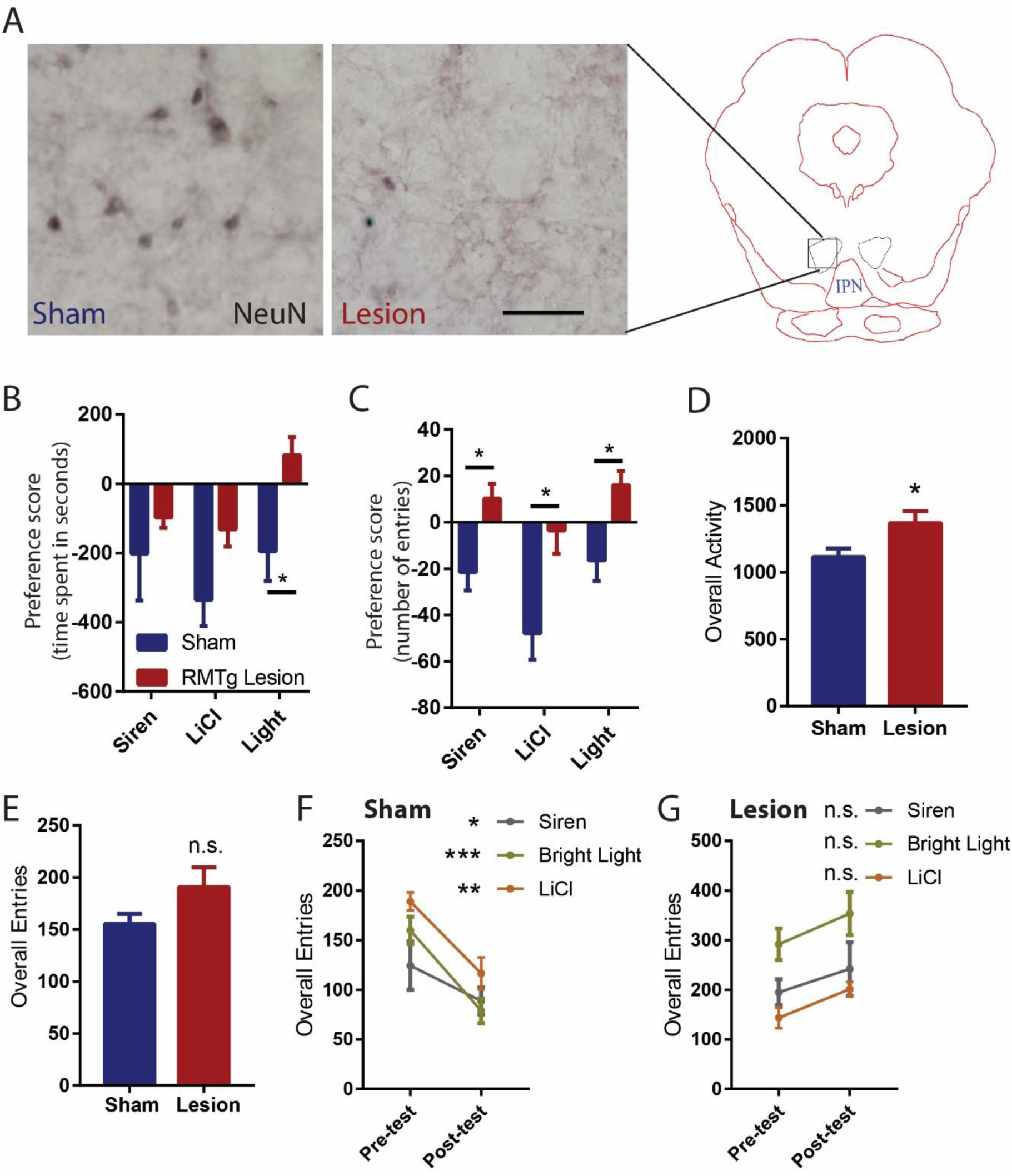
RMTg lesions disrupt conditioned place aversion to a wide range of stimuli. (**A**) Photographs of immunostaining of NeuN in RMTg region with and without excitotoxic lesions. Scalebar: 100um. (**B**) RMTg-lesioned rats showed smaller biases away from the stimulus-paired chambers. (**C**) RMTg lesions produced a somewhat stronger effect on the preference score as measured by entries. (**D**) RMTg lesioned animals showed slightly higher locomotor activity in conditioning chambers. (**E**) Overall entries to either chamber were unaffected by RMTg lesions. Total combined entries to both chambers were reduced after conditioning in all stimulus groups for sham rats (**F**), but not lesioned rats (**G**).

### Discussion

We show that in addition to being inhibited by reward-predictive cues, that RMTg neurons are activated by six distinct aversive stimuli of widely varying sensory modalities and timescales, while also playing key roles in driving VTA inhibitions and behavioral avoidance to aversive stimuli. Furthermore, we found evidence that removal of sustained aversive stimuli (LiCl and restraint stress) produces reward-correlated responses in the RMTg, while removal of a sustained reward (cocaine) produces aversion-correlated responses in the RMTg. These RMTg responses parallel actual reward and aversion as measured by conditioned place preference, and suggest possible neural substrates of the opponent process model proposed decades ago to describe the phenomenon that removal of a strong affective stimulus often produces an affective state in the opposite direction of the initial stimulus (Solomon and Corbit, 1973).

#### Broad encoding of valence in RMTg contributes to VTA valence encoding

The current study established a broad role for the RMTg in driving VTA inhibitions to aversive stimuli. Expect for footshocks and shock cues (Jhou et al., 2009), RMTg responses to other aversive stimulus modalities had been largely untested, with the exception of one study showing that many aversive stimuli do *not* induce c-Fos activation in the RMTg (Sanchez-Catalan et al., 2017). Additionally, previous studies had also identified several brain regions other than the RMTg that might also contribute to DA responses to aversive stimuli including ventral palladium and hypothalamus (Nieh et al., 2016; Tian et al., 2016; Tian and Uchida, 2015). Given the uncertainty of RMTg generalizability to a wide range of aversive stimuli and it influence on DA responses, it is notable that we saw RMTg activations by all six aversive stimuli that we tested (including several that did not induce c-Fos in the earlier study of Sanchez-Catalan et al.), and that RMTg lesions greatly reduced aversion-induced inhibitions to all tested stimuli in either classified pDA neurons or the entire VTA population.

Although our recorded pDA neurons were not genetically verified to be DAergic (e.g. by optogenetic phototagging), we used a classification method involving responses to reward-predictive cues that in mice is highly correlated with optogenetically identified DA neurons (Cohen et al., 2012; Matsumoto et al., 2016; Tian and Uchida, 2015), and that had been shown to be more accurate than traditional methods identifying DA neurons via long spike waveform durations and low firing rates (Cohen et al., 2012). Moreover, we found that aversion-induced inhibitory patterns were enriched in pDA neurons compared to all recorded VTA neurons, consistent with previous studies showing that DAergic neuron is the predominant cell-type exhibiting inhibitory responses to aversive stimuli in the VTA (Brischoux et al., 2009; Cohen et al., 2012; Ungless et al., 2004). Hence, we speculate that a majority of recorded pDA neurons are DAergic. However, because we noted a loss of aversive stimulus-induced inhibition across all VTA neurons, whether or not they had been classified as pDA, our findings that VTA encoding of aversive stimuli is RMTg-dependent does not rely on the ability to classify VTA neurons into pDA or non-pDA neurons.

#### Heterogeneity and opponency of RMTg firing patterns

In RMTg neurons, negative valence-encoding patterns, i.e. activation by aversive stimuli and inhibition by rewarding stimuli, were the most common, but we also found other response patterns in RMTg neurons. However, valence-encoding patterns were particularly enriched in VTA-projecting but not DRN-projecting neurons, as shown by endoscopic calcium imaging in mice. Thus, our results suggest that despite the heterogeneity in RMTg response patterns, that its influence on the VTA encoding of aversive stimuli is likely to be somewhat more uniform, and consistent with the originally proposed RMTg role in driving VTA inhibitions to aversive stimuli (Hong et al., 2011). Although calcium transients measured in the current study may not be strictly comparable to electrophysiological firing (Grewe et al., 2010; Jennings et al., 2015), calcium transients did show clear differences between VTA- and DRN-projecting neurons, strongly indicating that heterogeneity in RMTg activation patterns are correlated with differences in RMTg projection targets. However in contrast to the valence encoding roles of the VTA-projecting RMTg population, the roles of the DRN-projecting neurons are less clear, although we found that a high percentage of DRN-projecting RMTg neurons are activated by footshock, raising the possibility that the RMTg may also drive DRN inhibition by aversive stimuli such as footshock (Schweimer and Ungless, 2010).

In addition to RMTg activation to aversive stimuli and its influence on VTA responses to these stimuli, we also showed evidence that RMTg neurons could dynamically encode opponent processing of motivational states. RMTg responses to all three sustained stimuli (LiCl, restraint stress, and cocaine) exhibit a delayed “rebound” shift in the opposite direction to the initial responses. The RMTg opponent-firing responses were coincident with the opponent motivational states induced by the stimulus, suggesting parallels with opponent process theory that has been found in responses to many motivational stimuli across species (Becerra et al., 2013; Ettenberg, 2004; Jhou et al., 2013; Koob et al., 1989; Navratilova and Porreca, 2014; Navratilova et al., 2012; Solomon, 1980; Solomon and Corbit, 1973). Furthermore, previous studies have indicated an important DA role in opponent processing, as relief of pain induced increased DAergic signaling in the nucleus accumbens, which could be blocked by intra-VTA infusion of GABA agonists, and inhibition of mesolimbic dopamine system results in diminished relief learning (Becerra et al., 2013; Mayer et al., 2018; Navratilova and Porreca, 2014; Navratilova et al., 2012). Our results suggest that the opponent processing signals could be potentially driven by the RMTg through inhibiting and disinhibiting VTA DA neurons (Bourdy et al., 2014; Jalabert et al., 2011; Lecca et al., 2011; Lecca et al., 2012).

#### Contrast between RMTg and VTA GABA neurons

Finally, the current study also provides evidence on segregated functions between RMTg and the closely adjacent VTA GABAergic neurons. This contrasts somewhat with prior studies showing similar functions, e.g. that they have similar inhibitory influences on DA neurons, and that activation of either RMTg or VTA GABAergic neurons induce behavioral avoidance (Jhou et al., 2009; Lammel et al., 2012; Stamatakis and Stuber, 2012; Tan et al., 2012). However, a series of brilliant studies from Uchida’s group have recorded genetically identified GABAergic neurons in the VTA and show that they exhibit ramping activities in responses to reward cues and that they remain activated until animals receive rewards (Cohen et al., 2012; Eshel et al., 2015). In contrast, we showed that RMTg neurons were inhibited by reward cues and then quickly returned to baseline firing levels, a completely different firing pattern from VTA interneurons. Although Uchida’s studies are in mice, we also found that about 1/3 of our recorded VTA neurons exhibited similar ramping activities in responses to reward cues, a response pattern almost never seen in the RMTg recordings. Furthermore, only 6% (4/66) of VTA neurons exhibited negative valence encoding, the predominant response pattern seen in the RMTg. Furthermore, optogenetic inhibition of VTA GABA interneurons disrupts phasic DA responses to reward cues and results in a sustained DA activation throughout the entire presentation of cues (Eshel et al., 2015). In the current study, we showed that RMTg lesions removed majority of aversion-induced inhibition in VTA neurons, but did not alter the proportion nor responses of VTA neurons to reward cues. Therefore, our results suggest that RMTg neurons convey completely different information onto DA neurons than VTA GABAergic neurons, and potentially mediate distinct aspects of motivated behaviors.

These electrophysiological findings, taken together with our behavioral effects on conditioned place test, show a broad RMTg role in processing aversive stimuli. The RMTg is activated by aversive stimuli of a remarkably wide range of modalities and timescales, is a predominant driver of VTA inhibition by such aversive stimuli, and also drives place aversion to these stimuli.

## Methods

### Animals

All procedures were conducted under the National Institutes of Health Guide for the Care and Use of Laboratory Animals, and all protocols were approved by Medical University of South Carolina Institutional Animal Care and Use Committee. Adult male Sprague Dawley rats weighing 250 to 450 g from Charles River Laboratories were paired housed in standard shoebox cages with food and water provided ad libitum until experiments started. Rats were single housed during all experiments. In total, 64 rats were used for these experiments. 47 rats completed conditioned place test, of these, 24 rats had RMTg lesions. 15 rats underwent RMTg recordings, and 8 underwent VTA recordings.

### Surgeries

All surgeries were conducted under aseptic conditions with rats that were under isoflurane (1– 2% at 0.5–1.0 liter/min) anesthesia. Analgesic (ketoprofen, 5mg/kg) was administered subcutaneously immediately after surgery. Rats were given at least 5 days to recover from surgery. For recording experiments, drivable electrode arrays were implanted above the RMTg (AP: -7.4mm; ML: 2.1mm; DV: - 7.4mm from dura, 10-degree angle) or the VTA (AP: -5.5mm; ML: 2.5mm; DV: -7.8mm from dura, 10-degree angle). For lesion experiments, 50nl of 400mM quinolinic acid per side was injected with a glass pipette into the RMTg (AP: -7.6mm; ML: 2.1mm; DV: -7.8mm from dura, 10-degree angle). Sham controls received a saline infusion of equal volume into the RMTg. Rats were kept anesthetized with pentobarbital intraperitoneally (55mg/kg) for up to 3 hours’ post-surgery to reduce excitotoxic effect.

For intravenous catheterization, rats were implanted with 0.037-inch diameter silicone tubing into the jugular vein or the femoral vein. Intravenous lines were exteriorized at the back of the neck and flushed every 2 days with sterile saline and 0.1ml TCS lock solution to ensure patency.

Mice surgeries are described in the “calcium imaging” section.

### Perfusions and tissue sectioning

Rats used for all experiments were sacrificed with an overdose of isoflurane and perfused transcardially with 10% formalin in 0.1M phosophate buffered saline (PBS), pH 7.4. Brains from electrophysiology experiments had passage of 100μA current before perfusion, allowing electrode tips to be visualized. Brains were removed from the skull, equilibrated in 20% sucrose solution until sunk, and cut into 40μm sections on a freezing microtome. Sections were stored in phosphate buffered saline with 0.05% sodium azide.

### Immunohistochemistry for TH and FOXP1

Free-floating sections were immunostained for TH or FOXP1 by overnight incubation in mouse anti-TH (Millipore, MAB-377, 1: 10,000 dilution) or rabbit anti-FOXP1 (Abcam, ab16645, 1: 50,000 dilution) primary in PBS with 0.25% Triton-X and 0.05% sodium azide. Afterwards, tissue was washed three times in PBS and incubated in biotinylated donkey-anti-mouse or anti-rabbit secondary (1:1000 dilution, Jackson Immunoresearch, West Grove, PA) for 30 min, followed by three 30s rinses in PBS, followed by 1 hour in avidin-biotin complex (Vector). For TH-staining, tissue was then rinsed in sodium acetate buffer (0.1M, pH 7.4), followed by incubation for 5 min in 1% diaminobenzidine (DAB). For FOXP1 staining, nickel and hydrogen peroxide (Vector) were added to reveal a blue-black reaction product.

For florescent staining of FOXP1, free-floating sections were incubated in rabbit anti-FOXP1 (Abcam, ab16645, 1: 50,000 dilution) primary in PBS with 0.25% Triton-X and 0.05% sodium azide. Afterwards, tissue was washed three times in PBS and incubated in cy3-conjugated donkey-anti-rabbit secondary (1:1000 dilution, Jackson Immunoresearch, West Grove, PA).

### Behavioral training for electrophysiological recordings

Rats were food restricted to 85% of their *ad libitum* body weight and trained to associate distinct auditory cues with either a food pellet or no outcome. Behavior was conducted in standard Med Associates chambers (St. Albans, VT). Food-predictive and neutral tones were a 1 kHz tone (75 dB) and white noise (75dB), respectively. The food-predictive cue was presented for 2s, and a food pellet (45 mg, BioServ) was delivered immediately after cue offset. The neutral tone was also presented for 2s, but no food pellet was delivered. The two trial types were randomly presented with a 30s interval between successive trials. A “correct” response was scored if the animal either entered the food tray within 2 s after reward cues, or withheld a response for 2 s after neutral tones. Rats were trained with 100 trials per session, one session per day, until they achieved 85% accuracy in any 20-trial block. Once 85% accuracy was established, rats underwent surgeries. After recovery from surgeries, rats were then trained with one extra session in which neutral tone trials were replaced by aversive trials consisting of a 2s 8kHz tone (75dB) followed by a 10ms 0.7mA footshock.

### Electrophysiological recordings

After final training, electrodes consisted of a bundle of sixteen 18 μm Formvar-insulated nichrome wires (A-M system) attached to a custom-machined circuit board. Electrodes were grounded through a 37-gauge wire attached to a gold-plated pin (Newark Electronics), which was implanted into the overlying cortex. Recordings were performed during once-daily sessions, and electrodes were advanced 80-160 μm at the end of each session. The recording apparatus consisted of a unity gain headstage (Neurosys LLC) whose output was fed to preamplifiers with high-pass and low-pass filter cutoffs of 300 Hz and 6 kHz, respectively. Analog signals were converted to 18-bit values at a frequency of 15.625 kHz using a PCI card (National Instruments) controlled by customized acquisition software (Neurosys LLC). Spikes were initially detected via thresholding to remove signals less than twofold above background noise levels, and signals were further processed using principal component analysis performed by NeuroSorter software. Spikes were accepted only if they had a refractory period, determined by <0.2% of spikes occurring within 1ms of a previous spike, as well as by the presence of a large central notch in the auto-correlogram. Neurons that had significant drifts in firing rates were excluded. Since the shock duration used in the current study was 10ms, the first 10ms of data after footshock were removed in order to reduce shock artifacts.

For phasic aversive stimuli paradigm, rats were again placed on mild food deprivation, and recordings obtained in sessions consisting of 50 reward trials, followed by 4 different phasic aversive stimuli (10ms 0.7mA footshock, 2s 1600 lumens bright light presentation and 1s acoustic 115dB siren) randomly interleaved with 30s interval, followed by a 15 min baseline recording period, and then either 10 mg/kg LiCl or saline, i.p. or 6 min restraint stress. For VTA recordings, rats only completed the sessions with reward and phasic aversive stimuli. For Pavlovian conditioning paradigm, once rats achieved 85% accuracy in reward trials, they were trained to respond to an 8kHz tone (75dB) lasting for 2s followed by a mild footshock (0.7mA). During testing, rats again placed on mild food deprivation, and recordings obtained in sessions consisting of 150 mixture of reward trials, neutral trials, and shock trials randomly selected. Rats were recorded for one or two session per day, and electrodes advanced 80–160 μm at the end of each session. Neurons with significant reductions in baseline firing rates across sessions were excluded from the study, as this is indicative of drifting of recording wires between sessions.

### In vivo Ca^2+^ imaging

Wild type mice were placed into operant chambers (Med Associates) after being food deprived to 85% their original body weight. Mice were trained with 1 kHz auditory tones (70 dB, 2 seconds) followed immediately by sucrose pellet delivery. During training, tones came to elicit approach to the food tray. Once mice reached criterion (approach responses within 3 seconds on >85% of trials), ad libitum feeding was restored, and mice received injections of AAV2-hSyn-FLEX-CaMP6f virus (UNC Vector Core) into the RMTg (AP: -4; ML: 1.2, DV: -4 from dura, 10-degree angle) and CAV2-Cre (Montpellier Vector Core) into the VTA (AP: -5.1 mm; ML: 2.5mm; DV: -7.8mm from dura, 10-degree angle) or DRN (AP: -3.7 mm; ML: -0.6mm; DV: 3mm from dura, 10-degree angle). After 3 weeks’ recovery, we implanted a gradient index (GRIN) lens (outer diameter 0.5 mm, length 6.0 mm) (Inscopix) with the tip of the lens placed 0.2-0.3 mm dorsal to the viral injection site. Four to 6 weeks after lens implantation, a baseplate was implanted. Before each recording session, the mouse was briefly anesthetized with isoflurane and a miniature camera attached to the baseplate with a setscrew. Each animal was then put back into its home cage for 20 min to recover. We recorded from each animal for 2 sessions per day: reward session and shock session. In reward sessions, the mice received 30 2 second auditory cues paired with sucrose pellet deliveries. The interval between trials was 30s. In shock sessions, mice received 30 0.2mA footshocks at 30s intervals. The endoscopic camera was turned on for 20 second windows centered at the onset of reward-predictive cue or footshock. Ca^2+^ images were acquired at 20 Hz at LED intensity from 30-70% and gain from 1-3.5. Using Mosaic software (Inscopix) we then spatially down-sampled videos to 400×384 pixels, and temporally down-sampled to 5 Hz. Individual trials (for reward/shock sessions) were concatenated and motion corrected. Calcium signals (dF/F) from individual cells were extracted by by CNMF-E, which extracts cellular calcium signals with minimal influence from the background (Pnevmatikakis et al., 2016). For reward/shock sessions, time courses were calculated for each 20-second interval corresponding to individual trials, and then averaged together for the session. Data points were normalized to baseline calcium activity, and the data were then smoothed using a 0.6-second moving average.

### Conditioned place test

Two groups of rats were tested for place conditioning after recovery from surgery. One group received bilateral RMTg lesions and the other group served as sham controls. The lesion was made prior to behavioral training, and lesion sizes were verified by NeuN staining. Both groups were exposed to intraperitoneal lithium chloride (150mg/kg, Sigma-Aldrich, 7 lesions and 6 shams), bright light (1600 lumens, 2s pulse with 2s duration, 7 lesions and 8 shams) and siren (115dB, 1s pulse with 2s duration, 7 lesions and 6 shams). We used i.p. saline injection, dim house light and 75dB noise as control treatments. The conditioned place test experiments were performed using a three-chambered apparatus (Med Associates, St Albans, VT) under dim room light. Each animal received only one aversive treatment or control treatment. On the first day of each experiment, rats explored all three chambers freely for 15 min once per day, and an average baseline preference score was determined for each. Over the next 8 days, rats performed one aversive session and one control session each day. Rats were placed into their previously preferred chambers for 15 min in aversive sessions and into the opposite chambers in control sessions. The order of treatments was counterbalanced across rats. On the ninth day, rats again explored all chambers freely without stimulus exposure. The preference score was defined as the number of seconds spent or number of entries into the stimulus-paired chamber minus the number of seconds spent in the unpaired chamber. We also calculated each animal’s preference shift, defined as the post-training preference minus the pre-training preference score. The overall activity of each animal was measured as the number of photobeam breaks.

### Statistical analysis of electrophysiological and behavioral data

RMTg and VTA neuron firing rates in response to phasic stimuli were calculated in 50ms bins and normalized to 1-second baseline before the onset of stimuli. RMTg firing rates in response to sustained stimuli were calculated in 5-minute bins and normalized to 15-minute baseline. Neurons with large drifting of the microwire electrodes during recordings were excluded from further analysis. Electrophysiology data were first tested for normality, then transformed to ranked forms if data failed tests of normality (p < 0.05, D’Agostino-Pearson test). Latency to maximum responses was calculated as the first bin that reached the peak or trough. Burst analysis was performed with NeuroExplorer, with a burst defined by any pair of spikes less than 80ms apart, and continuing until the last spike is more than 160ms separated from the following spike.

Significant responses in neural firing were determined by a threshold of p<0.05 for each neuron’s firing rate versus baseline (Wilcoxon signed-rank test for phasic stimuli trials in wire recording experiments, paired t-test for Ca2^+^ recording experiments). Calcium recording data and conditioned place aversion data passed tests of normality (p > 0.05, D’Agostino-Pearson test), and were analyzed using parametric tests. Post hoc tests after one-way and two-way ANOVA were Holm-Sidak and Bonferroni, respectively. Calculations were performed using Matlab (Mathworks) and Prism 7 software (Graph Pad).

## Author Contributions

H.L., D.P., J.Y.C. and M.E. conducted the experiments; H.L. designed the experiments and wrote the paper; H.L. and T.C.J. reviewed and edited the paper.

Conceptualization, H.L. and T.C.J.; Methodology, H.L. and T.C.J.; Investigation, H.L., D.P., J.Y.C. and M.E.; Writing – Original Draft, H.L.; Writing – Review & Editing, H.L. and T.C.J.; Funding Acquisition, T.C.J.

## Declaration of Interests

No conflicts in interests

**Supplementary Fig1.**
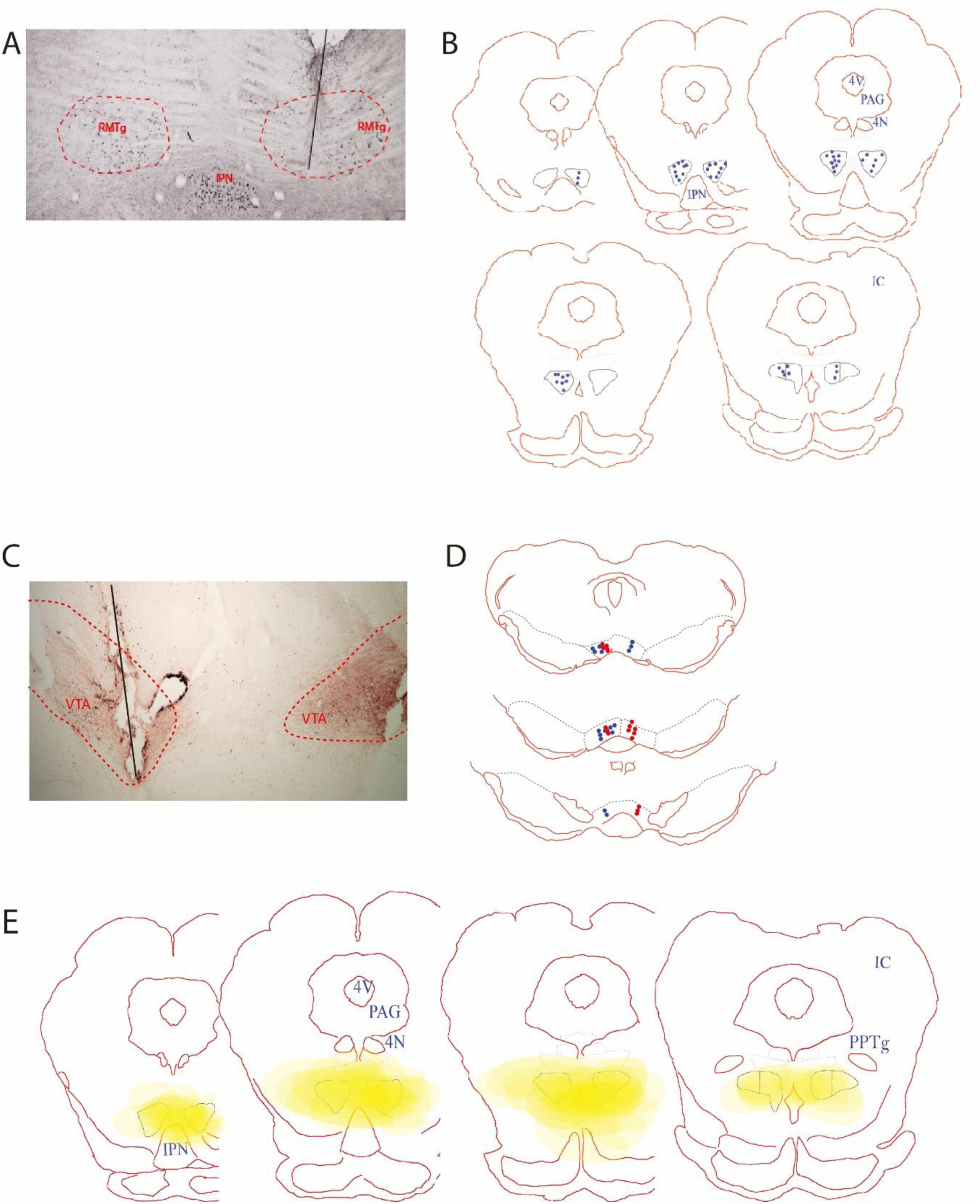
(**A**) Photo of an electrode trace in the RMTg region stained with FOXP1 (black dots). Red dashed line: the shape of the RMTg. Black line: electrode. (**B**) Recording sites in the RMTg region verified by FOXP1 staining. (**C**) Photo of an electrode trace in the RMTg region stained with TH. Red dashed line: the shape of the VTA. Black line: electrode. (**D**) Recording sites in the RMTg region verified by TH staining. Blue dots: intact rats. Red dots: lesion rats. (**E**) RMTg (black circles) lesion sizes (yellow) in conditioned place aversion experiments.

**Supplementary Fig2.**
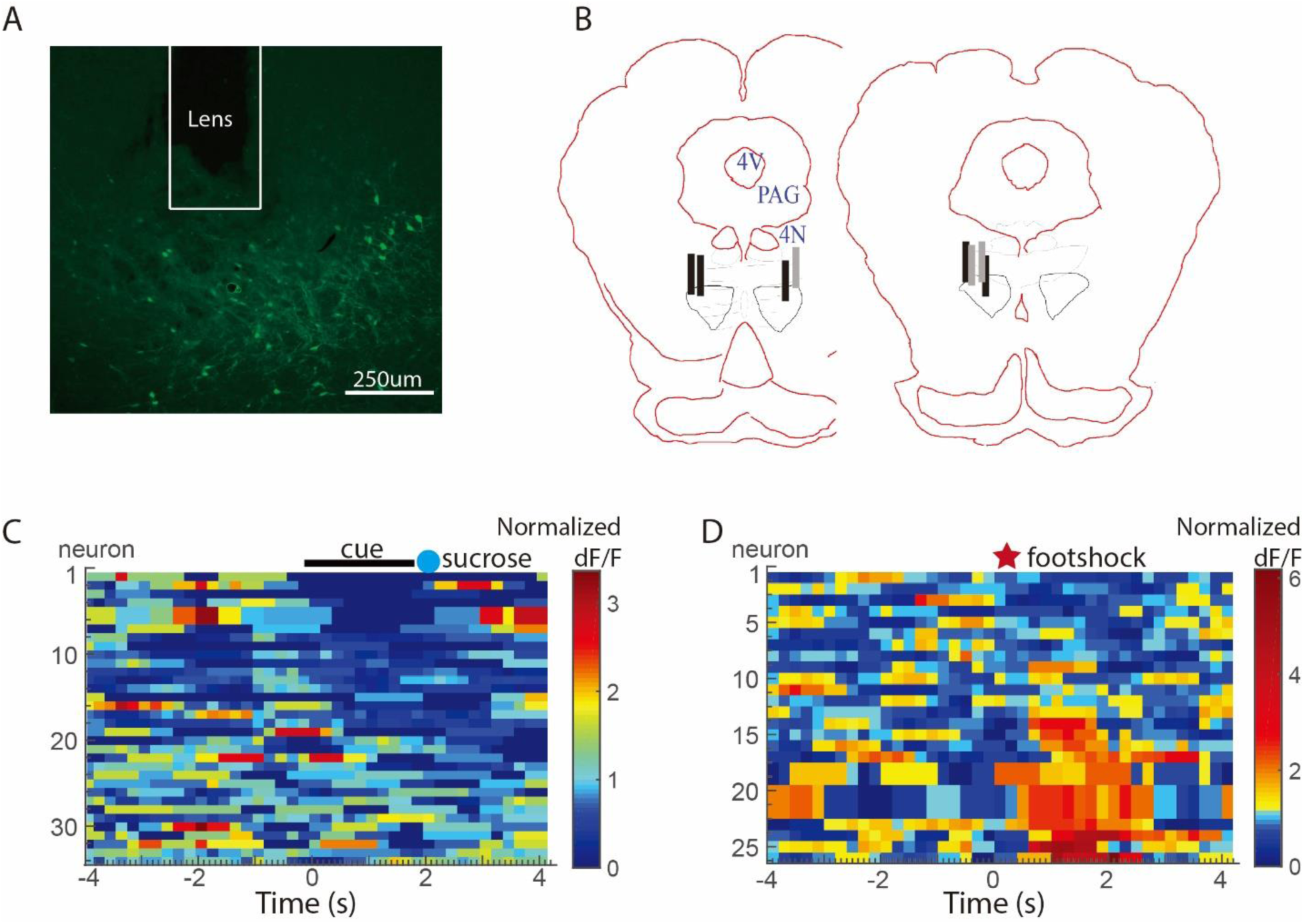
(**A**) Photo of a GRIN lens trace in the RMTg. Green cells are gCamP6 positive cells. (**B**) Lens placements (Black: VTA-projecting group; Gray: DRN-projecting group). (**C, D**) Calcium signals (dF/F) from individual VTA-projecting RMTg neurons in response to reward cues and footshocks. Black bar: duration of the cue; Blue circle: sucrose deliver; Red star: onset of footshocks.

**Supplementary Fig3.**
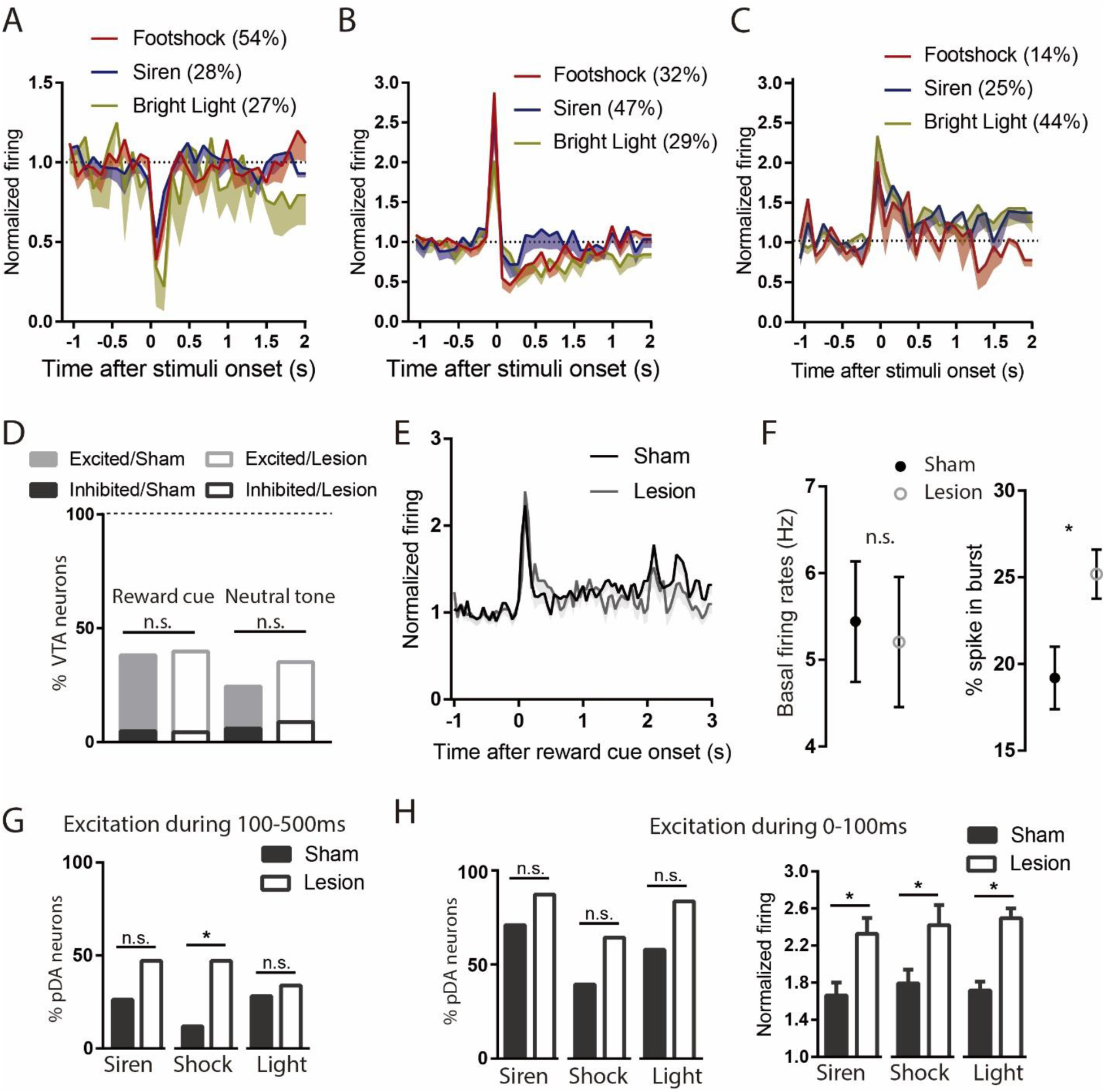
In RMTg intact animals, pDA neurons responded to aversive stimuli with pure inhibition (**A**), inhibition after a brief excitation (**B**), or pure excitation (**C**). The percentages of neurons responding to different aversive stimuli in each category are listed in each panel. (**D**) Percentages of pDA neurons with excitatory/inhibitory responses to reward/neutral tones were not affected by RMTg lesions. (**E**) RMTg lesions did not alter pDA responses to reward cue. (**F**) RMTg lesion did not change basal firings, but increased the percentage of pDA spikes in bursts. (**G**) RMTg lesion slightly increased proportion of pDA neurons excited by aversive stimuli during 100-500ms post-stimulus. (**H**) The percentages of pDA neurons with initial excitations to aversive stimuli were marginally increased after RMTg lesion, while the magnitudes of these initial excitations were significantly increased.

## Acknowledgements

This work was funded by National Institutes of Health grants R01 DA037327 and R21 DA032898, both to TCJ.

## References

Balcita-Pedicino, J.J., Omelchenko, N., Bell, R., and Sesack, S.R. (2011). The inhibitory influence of the lateral habenula on midbrain dopamine cells: ultrastructural evidence for indirect mediation via the rostromedial mesopontine tegmental nucleus. J Comp Neurol 519, 1143–1164.

Becerra, L., Navratilova, E., Porreca, F., and Borsook, D. (2013). Analogous responses in the nucleus accumbens and cingulate cortex to pain onset (aversion) and offset (relief) in rats and humans. J Neurophysiol 110, 1221–1226.

Bourdy, R., Sanchez-Catalan, M.J., Kaufling, J., Balcita-Pedicino, J.J., Freund-Mercier, M.J., Veinante, P., Sesack, S.R., Georges, F., and Barrot, M. (2014). Control of the nigrostriatal dopamine neuron activity and motor function by the tail of the ventral tegmental area. Neuropsychopharmacology 39, 2788–2798.

Brischoux, F., Chakraborty, S., Brierley, D.I., and Ungless, M.A. (2009). Phasic excitation of dopamine neurons in ventral VTA by noxious stimuli. Proc Natl Acad Sci U S A 106, 4894–4899.

Brown, R.M., Kupchik, Y.M., Spencer, S., Garcia-Keller, C., Spanswick, D.C., Lawrence, A.J., Simonds, S.E., Schwartz, D.J., Jordan, K.A., Jhou, T.C., et al. (2017). Addiction-like Synaptic Impairments in Diet-Induced Obesity. Biol Psychiatry 81, 797–806.

Chang, C.Y., Esber, G.R., Marrero-Garcia, Y., Yau, H.J., Bonci, A., and Schoenbaum, G. (2016). Brief optogenetic inhibition of dopamine neurons mimics endogenous negative reward prediction errors. Nat Neurosci 19, 111–116.

Cohen, J.Y., Haesler, S., Vong, L., Lowell, B.B., and Uchida, N. (2012). Neuron-type-specific signals for reward and punishment in the ventral tegmental area. Nature 482, 85–88.

Eshel, N., Bukwich, M., Rao, V., Hemmelder, V., Tian, J., and Uchida, N. (2015). Arithmetic and local circuitry underlying dopamine prediction errors. Nature 525, 243–246.

Ettenberg, A. (2004). Opponent process properties of self-administered cocaine. Neurosci Biobehav Rev 27, 721–728.

Ettenberg, A., Raven, M.A., Danluck, D.A., and Necessary, B.D. (1999). Evidence for opponent-process actions of intravenous cocaine. Pharmacol Biochem Behav 64, 507–512.

Fiorillo, C.D., Song, M.R., and Yun, S.R. (2013). Multiphasic temporal dynamics in responses of midbrain dopamine neurons to appetitive and aversive stimuli. J Neurosci 33, 4710–4725.

Grewe, B.F., Langer, D., Kasper, H., Kampa, B.M., and Helmchen, F. (2010). High-speed in vivo calcium imaging reveals neuronal network activity with near-millisecond precision. Nat Methods 7, 399–405.

Henny, P., Brown, M.T., Northrop, A., Faunes, M., Ungless, M.A., Magill, P.J., and Bolam, J.P. (2012). Structural correlates of heterogeneous in vivo activity of midbrain dopaminergic neurons. Nat Neurosci 15, 613–619.

Hong, S., Jhou, T.C., Smith, M., Saleem, K.S., and Hikosaka, O. (2011). Negative reward signals from the lateral habenula to dopamine neurons are mediated by rostromedial tegmental nucleus in primates. J Neurosci 31, 11457–11471.

Jalabert, M., Bourdy, R., Courtin, J., Veinante, P., Manzoni, O.J., Barrot, M., and Georges, F. (2011). Neuronal circuits underlying acute morphine action on dopamine neurons. Proc Natl Acad Sci U S A 108, 16446–16450.

Jennings, J.H., Sparta, D.R., Stamatakis, A.M., Ung, R.L., Pleil, K.E., Kash, T.L., and Stuber, G.D. (2013). Distinct extended amygdala circuits for divergent motivational states. Nature 496, 224–228.

Jennings, J.H., Ung, R.L., Resendez, S.L., Stamatakis, A.M., Taylor, J.G., Huang, J., Veleta, K., Kantak, P.A., Aita, M., Shilling-Scrivo, K., et al. (2015). Visualizing hypothalamic network dynamics for appetitive and consummatory behaviors. Cell 160, 516–527.

Jhou, T.C., Fields, H.L., Baxter, M.G., Saper, C.B., and Holland, P.C. (2009). The rostromedial tegmental nucleus (RMTg), a GABAergic afferent to midbrain dopamine neurons, encodes aversive stimuli and inhibits motor responses. Neuron 61, 786–800.

Jhou, T.C., Good, C.H., Rowley, C.S., Xu, S.P., Wang, H., Burnham, N.W., Hoffman, A.F., Lupica, C.R., and Ikemoto, S. (2013). Cocaine drives aversive conditioning via delayed activation of dopamine-responsive habenular and midbrain pathways. J Neurosci 33, 7501–7512.

Kaufling, J., Veinante, P., Pawlowski, S.A., Freund-Mercier, M.J., and Barrot, M. (2009). Afferents to the GABAergic tail of the ventral tegmental area in the rat. J Comp Neurol 513, 597–621.

Koob, G.F., Stinus, L., Le Moal, M., and Bloom, F.E. (1989). Opponent process theory of motivation: neurobiological evidence from studies of opiate dependence. Neurosci Biobehav Rev 13, 135–140.

Kovacs, K.J. (2008). Measurement of immediate-early gene activation- c-fos and beyond. J Neuroendocrinol 20, 665–672.

Lahti, L., Haugas, M., Tikker, L., Airavaara, M., Voutilainen, M.H., Anttila, J., Kumar, S., Inkinen, C., Salminen, M., and Partanen, J. (2016). Differentiation and molecular heterogeneity of inhibitory and excitatory neurons associated with midbrain dopaminergic nuclei. Development 143, 516–529.

Lammel, S., Lim, B.K., Ran, C., Huang, K.W., Betley, M.J., Tye, K.M., Deisseroth, K., and Malenka, R.C. (2012). Input-specific control of reward and aversion in the ventral tegmental area. Nature 491, 212– 217.

Lecca, S., Melis, M., Luchicchi, A., Ennas, M.G., Castelli, M.P., Muntoni, A.L., and Pistis, M. (2011). Effects of drugs of abuse on putative rostromedial tegmental neurons, inhibitory afferents to midbrain dopamine cells. Neuropsychopharmacology 36, 589–602.

Lecca, S., Melis, M., Luchicchi, A., Muntoni, A.L., and Pistis, M. (2012). Inhibitory inputs from rostromedial tegmental neurons regulate spontaneous activity of midbrain dopamine cells and their responses to drugs of abuse. Neuropsychopharmacology 37, 1164–1176.

Li, Y., Zhong, W., Wang, D., Feng, Q., Liu, Z., Zhou, J., Jia, C., Hu, F., Zeng, J., Guo, Q., et al. (2016). Serotonin neurons in the dorsal raphe nucleus encode reward signals. Nat Commun 7, 10503.

Matsumoto, H., Tian, J., Uchida, N., and Watabe-Uchida, M. (2016). Midbrain dopamine neurons signal aversion in a reward-context-dependent manner. Elife 5.

Mayer, D., Kahl, E., Uzuneser, T.C., and Fendt, M. (2018). Role of the mesolimbic dopamine system in relief learning. Neuropsychopharmacology 43, 1651–1659.

Navratilova, E., and Porreca, F. (2014). Reward and motivation in pain and pain relief. Nat Neurosci 17, 1304–1312.

Navratilova, E., Xie, J.Y., Okun, A., Qu, C., Eyde, N., Ci, S., Ossipov, M.H., King, T., Fields, H.L., and Porreca, F. (2012). Pain relief produces negative reinforcement through activation of mesolimbic reward-valuation circuitry. Proc Natl Acad Sci U S A 109, 20709–20713.

Nieh, E.H., Vander Weele, C.M., Matthews, G.A., Presbrey, K.N., Wichmann, R., Leppla, C.A., Izadmehr, E.M., and Tye, K.M. (2016). Inhibitory Input from the Lateral Hypothalamus to the Ventral Tegmental Area Disinhibits Dopamine Neurons and Promotes Behavioral Activation. Neuron 90, 1286–1298.

Pnevmatikakis, E.A., Soudry, D., Gao, Y., Machado, T.A., Merel, J., Pfau, D., Reardon, T., Mu, Y., Lacefield, C., Yang, W., et al. (2016). Simultaneous Denoising, Deconvolution, and Demixing of Calcium Imaging Data. Neuron 89, 285–299.

Sanchez-Catalan, M.J., Faivre, F., Yalcin, I., Muller, M.A., Massotte, D., Majchrzak, M., and Barrot, M. (2017). Response of the Tail of the Ventral Tegmental Area to Aversive Stimuli. Neuropsychopharmacology 42, 638–648.

Schweimer, J.V., and Ungless, M.A. (2010). Phasic responses in dorsal raphe serotonin neurons to noxious stimuli. Neuroscience 171, 1209–1215.

Solomon, R.L. (1980). The opponent-process theory of acquired motivation: the costs of pleasure and the benefits of pain. Am Psychol 35, 691–712.

Solomon, R.L., and Corbit, J.D. (1973). An opponent-process theory of motivation. II. Cigarette addiction. J Abnorm Psychol 81, 158–171.

Stamatakis, A.M., and Stuber, G.D. (2012). Activation of lateral habenula inputs to the ventral midbrain promotes behavioral avoidance. Nat Neurosci 15, 1105–1107.

Tan, K.R., Yvon, C., Turiault, M., Mirzabekov, J.J., Doehner, J., Labouebe, G., Deisseroth, K., Tye, K.M., and Luscher, C. (2012). GABA neurons of the VTA drive conditioned place aversion. Neuron 73, 1173– 1183.

Tian, J., Huang, R., Cohen, J.Y., Osakada, F., Kobak, D., Machens, C.K., Callaway, E.M., Uchida, N., and Watabe-Uchida, M. (2016). Distributed and Mixed Information in Monosynaptic Inputs to Dopamine Neurons. Neuron 91, 1374–1389.

Tian, J., and Uchida, N. (2015). Habenula Lesions Reveal that Multiple Mechanisms Underlie Dopamine Prediction Errors. Neuron 87, 1304–1316.

Tomasiewicz, H.C., Mague, S.D., Cohen, B.M., and Carlezon, W.A., Jr. (2006). Behavioral effects of short-term administration of lithium and valproic acid in rats. Brain Res 1093, 83–94.

Tzschentke, T.M. (2007). Measuring reward with the conditioned place preference (CPP) paradigm: update of the last decade. Addict Biol 12, 227–462.

Ungless, M.A., Magill, P.J., and Bolam, J.P. (2004). Uniform inhibition of dopamine neurons in the ventral tegmental area by aversive stimuli. Science 303, 2040–2042.

Vento, P.J., Burnham, N.W., Rowley, C.S., and Jhou, T.C. (2017). Learning From One’s Mistakes: A Dual Role for the Rostromedial Tegmental Nucleus in the Encoding and Expression of Punished Reward Seeking. Biol Psychiatry 81, 1041–1049.

Winston, C.R., Leavell, B.J., Ardayfio, P.A., Beard, C., and Commissaris, R.L. (2001). A nonextinction procedure for long-term studies of classically conditioned enhancement of acoustic startle in the rat. Physiol Behav 73, 9–17.

